# Possible origin of the scrapie genome in small endogenous RNAs; studies on eight candidate species in 263K scrapie-infected hamster brain

**DOI:** 10.1101/620732

**Authors:** Eleanor Barnard, Kathryn Estibeiro, Rory Duncan, Janet Baird, David Fettes, Jacqueline Wood, Hugh Fraser, Peter Estibeiro, Richard Lathe

**Affiliations:** Centre for Genome Research and Centre for Neuroscience, University of Edinburgh, King’s Buildings, West Mains Road, Edinburgh EH9 3JQ, UK; Institute for Animal Health, Neuropathogenesis Unit, Ogston Building, West Mains Road, Edinburgh EH9 3JF, UK

## Abstract

The identity of the etiologic agent of the transmissible spongiform encephalopathies (TSEs), including bovine spongiform encephalopathy (BSE), scrapie and Creutzfeldt-Jakob disease (CJD), remains unknown. While much attention has been given to the hypothesis that the TSEs may be caused by a proteinaceous infectious agent or ‘prion’, there is considerable evidence to suggest that this hypothesis is incomplete. We have pursued an alternative contention: that the etiologic agent comprises in part a modified and replicating form of an endogenous nucleic acid, probably RNA. The ‘endovirus’ hypothesis contends that the parental molecule is most likely to be a small and highly-structured cellular RNA that can convert to a replicating molecule by a finite number of nucleotide sequence changes. We have begun a systematic analysis of candidate molecular species present in hamster brain infected with scrapie strain 263K. Initial work focussed on the 7S group of small RNAs. Examination of 7-2, 7SK and 7SL failed to reveal differences in abundance and/or sequence between normal and scrapie (263K)-infected hamster brain. Inspection of other possible candidates, including U3, H1/8-2 and novel molecules KR1, nu1 and nu2, similarly failed to provide evidence for scrapie-specific molecular variants; alterations to the KR1 sequence failed to correlate with disease. We present sequences of hamster RNAs 7-2, 7SK, 7SL, H1/8-2, U3, nu1, nu2 and KR1. Together our data so far fail to contradict or confirm the hypothesis, while arguing that the major species of these 8 RNA molecules are unlikely to correspond to the etiologic agent of the TSEs.

## INTRODUCTION

The transmissible encephalopathies (TSEs), including scrapie in sheep, bovine spongiform encephalopathy (BSE) in cattle and Kuru, Creutzfeldt-Jakob Disease (CJD) and Gerstmann-Sträussler-Scheinker syndrome (GSS) in human, are caused by unknown agents. Infection leads to progressive neurological disease and death. Diseased brain often exhibits a distinctive spongiform degeneration with amyloid deposition of an aggregated and protease-resistant form of a normal host protein, PrP^sc^, often referred to as the ‘prion’ protein, that is encoded by the *PRN-P* gene in human and the *Prn-p* gene in mouse (reviewed by Stahl & Prusiner, 1991; Prusiner & DeArmond, 1994).

The TSEs are particularly unusual in that they exhibit both a genetic and a transmissible component. Kuru was shown 30 years ago to be transmissible to chimpanzees (Gajdusek et al., 1966) while CJD can be transmitted to rodents by inoculation of infectious material (Manuelidis et al., 1978; Tateishi et al., 1978). However, some CJD cases, termed sporadic, appear to occur in the absence of any known contact with an infectious agent, suggesting that the disease may arise spontaneously. In addition, the genotype at the locus encoding the PrP protein clearly modulates susceptibility and/or onset of disease. Different alleles of the genes encoding PrP in mouse and sheep are associated with different incubation periods of the disease (Dickinson et al., 1968; Dickinson and Meikle, 1971; Carlson et al., 1986;1988; Hunter et al., 1989; Goldmann et al., 1990; 1991). Further, familial CJD/GSS is associated with mutations in the *PRN-P* gene (reviewed by Prusiner & Hsiao, 1994); the involvement of *PRN-P* mutations and polymorphisms in CJD in the UK is described by Windl et al. (1996).

Although the identity of the agent is unknown, infectivity appears refractory to treatments designed to eliminate nucleic acids (Alper et al., 1967, 1978; Latarjet et al., 1970) and attempts to identify a nucleic acid genome by a variety of approaches have so far failed (reviewed by Aiken and Marsh, 1990). Infectivity is also sensitive to detergents and to denaturation with phenol or chaotropic agents (Prusiner et al., 1980, 1981a, 1981b), and infectivity copurifies in some systems with PrP^sc^, leading to the protein-only hypothesis (Alper et al., 1967, 1978; Griffith, 1967; Prusiner et al., 1982). Under this hypothesis the agent is comprised principally of abnormal forms of PrP protein such as PrP^sc^; propagation takes place by recruitment of host PrP protein to these abnormal forms and the modified PrP protein hence constitutes the infectious agent (Prusiner, 1991). However, it is unclear whether the PrP^sc^ molecule itself could comprise the agent, for serial dilution experiments indicate that the minimal infectious unit equates to some 10^4^ copies of PrP^sc^ (Scott et al., 1991).

The ‘prion’ hypothesis also fails to explain the diversity of known strains of the agent that differ in incubation period, type of pathology, and end-point titre (Kimberlin et al., 1989; Fraser, 1993). For example, the properties of BSE agent are well-conserved on passage through several different mammalian species despite significant differences in the sequences of their PrP proteins (Bruce et al., 1997). Allotypic interactions and differential folding or modification of the PrP polypeptide have been invoked as a possible explanation for this conservation (Carlson et al., 1994). The recent report of neurodegenerative disease induced in mice by inoculation of BSE material, but in the absence of detectable PrP^sc^ (Lamézas et al., 1997), provides further strong evidence against the protein-only or prion hypothesis.

The agent is also sensitive to alkali (Prusiner et al., 1981a), suggestive of an RNA component. Radiation inactivation and other data have also been held to be consistent with the existence of a small scrapie-specific genome (Diringer and Kimberlin, 1983; Rohwer, 1984; 1986; Dees et al., 1985) and nucleic acids of substantial length are found in conventional preparations of PrP^sc^ (Kellings et al., 1992). Indeed, the involvement of conventional retroviral (Manuelidis et al., 1988) or even bacterial (Bastian, 1993) etiologic agents has been suggested. Small DNAs (eg. Aiken et al., 1990), RNAs (Akowitz et al., 1994) and virus-like particles (Ozel & Diringer, 1994) have been reported in scrapie-infected brain or in partially purified infectious material; these reports are so far unconfirmed.

The co-prion hypothesis (Weissmann, 1991) was proposed to explain the unusual properties of the agent and suggests that a suitably-modified PrP polypeptide comprises the basal infectious agent (prion); this complexes with a small nucleic acid species (co-prion) that modifies the properties of the agent to generate different strains. The hypothesis predicts that intense irradiation of different TSE strains should eliminate both the nucleic acid species and strain differences, residual infectivity remaining unaffected. This prediction has not been confirmed (HF, unpublished observations).

In an attempt to reconcile all the known properties of the agent we suggest that the agent comprises, at least in part, a structurally-modified and replicating version(s) of a small host-encoded (or otherwise environmentally abundant) nucleic acid(s). Our hypothesis, presented in Figure 1, contends that agent may be generated sporadically by mutations or other alterations of cellular RNAs (or genes encoding them) that lead to replication and propagation of the RNA molecule (possibly facilitated by PrP protein; Discussion). We term this contention the ‘endovirus’ hypothesis. The modification of an endogenous RNA to a disease-causing molecule would explain why subtractive hybridization procedures have so far failed to reveal a scrapie-specific nucleic acid (Oesch et al., 1988), though some transcripts are clearly upregulated in scrapie infection (Duguid et al., 1989).

**Figure 1.**
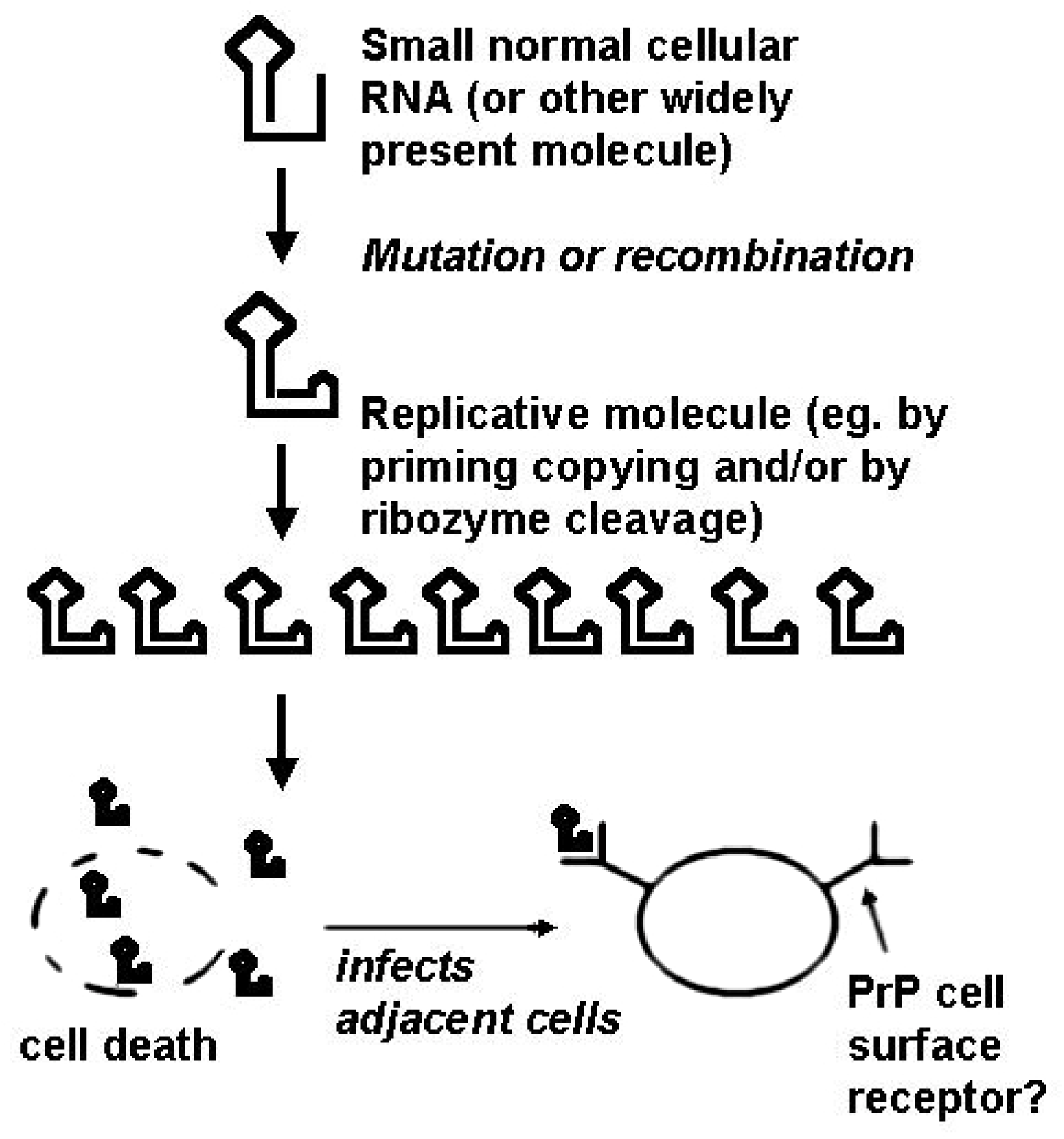
Schematic representation of the ‘endovirus’ hypothesis (text for details).

The agent seems unlikely to comprise a free RNA, however, because purified RNA from scrapie-infected mouse brain failed to transmit infectivity (Hunter et al., 1976). Nonetheless, in this study the presence of the RNA in a form that is poorly extracted by conventional procedures (for instance, a highly stable protein/RNA or even lipid/RNA complex) could not be ruled out.

RNA components of small nuclear or cytoplasmic ribonucleoproteins are possible candidate parental molecules: they are abundant, form tight complexes with other cellular components, and their high degree of secondary structure may predispose to replicative processes. They also play key roles in cellular housekeeping and metabolism: overabundance or dysfunction of such a molecule could disrupt such processes and cause neurodegeneration. The endovirus hypothesis favours replication by aberrant copying of RNA by endogenous polymerases, but does not exclude other mechanisms of replication, for instance by retrointegration into the genome and transcriptional amplification of the modified RNA. Indeed, pseudogenes of the small cellular RNAs U3 and 7SK appear to arise through such a process (Bernstein et al., 1983; Suh et al., 1989). In this context it is notable that some plant viroid RNAs contain regions of similarity with cellular RNAs (Haas et al., 1988), while in at least some instances satellite RNAs (virusoids) of plant viruses appear to arise spontaneously, presumably by alteration of endogenous sequences (reviewed by Roossinck et al., 1992).

We therefore set out to examine candidate molecular species in the brain of scrapie-infected hamsters with a view to determining whether any might differ in sequence or abundance between infected brain and controls.

## METHODS

### Brain samples and RNA preparation

Frozen brains from golden hamsters (LVG) either terminally infected with strain 263K scrapie, or control littermates, were prepared by the Neuropathogenesis Unit, Edinburgh. To purify RNA, brain samples were dispersed in RNAzol B, extracted with chloroform, centrifuged, and RNA precipitated from the supernatent with isopropanol and washed with 75% ethanol according to the recommendations of the RNAzol distributor (Biogenesis, Poole, UK). RNA was taken up in 1 mM MgCl_2_ and treated with DNase I (Pharmacia; 1 μg/ml, 2 h, 37°C) in the presence of RNAguard (950 units/ml) to eliminate DNA, reprecipitated as before and taken up in 0.5% sodium dodecyl sulfate (SDS).

### PCR primers

The PCR primers used in this work were: 7SL 5’ and 3’ primers were 5’-d*CACGAATTC*AGGCGCCGGGCGCGG-3’ and 5’-d*CACGAATTC*AGAGA CGGGGTCTCG-3’ respectively, designed according to the sequence of human 7SL RNA (HSRNA7SL); 7SK 5’ and 3’ primers were 5’-d*CACGAATTC* GGATGTGAGGCGATC-3’ and 5’-d*CACGAATTC*AAAGAAAGGCAGACT-3’, designed according to rat 7SK RNA sequence (RATSR7K); 7-2 primers were 5’-dAGCTCG<u>G</u>T<u>G</u>TGAAGG-3’ and 5’-dTAGCCGCGCTGAGAA, according to mouse 7-2 sequence (MUSMRP), 2 design errors in the 5’ primer (underlined) did not appear to affect amplification [see also below]; nu1-specific primers were 5’-dCAGGAGCCATCTGAGCCAAACAATCT-3’ and 5’-dTGAACTATTCTGGGACC AGTCATAGA-3’ (this work); nu2-specific primers were 5’-dTCAGAAGACA ATCTTCAGGGGCCAGT-3’ and 5’-dACACCAAAGCCCAGGGTCCTCCCA-3’ (this work), H1 primers were 5’-dGAGGGAAGCTCATCA-3’ and 5’-dAGAGTAGTCTGAATTGGGT-3’, designed according to a consensus of human and mouse H1/8-2 sequences (HSH1RNA and MUSRNASEP/MMU24680); U3 primers were 5’-dGACTATACTTT<u>G</u>AGGGAT-3’ (underlined, design error) and 5’-dACCACTCAGACCGCGTTC-3’, designed according to regions conserved between human (HUMUG3PE) and rat (RATUR3A) U3 sequences.

### PCR amplification, cloning and characterization

Reverse transcription and PCR (RTPCR) were performed as follows. 0.5 μg total RNA (DNase treated), 100 pMol 3’ primer, 2.5 mM MgCl_2_, 1 mM each dNTP, 30 units RNAguard (Pharmacia) and 12 units MMLV reverse transcriptase (Pharmacia) were incubated in a volume of 20 □ l at 23°C, 10 min; 40°C, 45 min, in 1 × PCR thermo DNA polymerase reaction buffer (Promega). Following the addition of 100 pMol each 5’ and 3’ primer and MgCl_2_ to 4 mM, final volume 50 μl, the mixture was incubated at 95°C, 5 min, and allowed to cool to room temperature. Amplification was commenced by addition of 3 units of ULTma DNA polymerase (Perkin-Elmer) in 50 μl ULTma buffer to give a final volume of 100 μl with a final concentration of 2 mM MgCl_2_; following preincubation (95°C, 5 min); 30 cycles (55°C, 30 sec; 72°C, 30 sec, 95°C, 30 sec) of amplification were performed, followed by a stop period (25°C, 1 min). Cloning of PCR products (either by restriction and ligation or by the T/A overlap technique), DNA sequencing and, Northern, Southern and colony hybridization were all according to standard procedures.

## RESULTS

### The etiologic agent and small cellular RNAs: candidate molecular species

To explore predictions made by the hypothesis we first examined the abundance and distribution of small nucleic acids, particularly RNAs. Gel electrophoresis on total nucleic acid from normal rodent brain and rat PC12 phaeochromocytoma cells revealed 20 or more small (100-1000 nt) abundant nucleic acid species; these were all RNA in virtue of their sensitivity to alkaline hydrolysis (not shown). No gross differences were observed when RNA preparations from normal and scrapie (263K)-infected hamster brain were compared (data not shown).

To refine our search we focused on the predicted replicative ability of the agent. Candidate mechanisms for replication include ribozyme-type cleavage (Pyle, 1993) of replicative intermediates, and/or self-priming of polymerization (see below). However, preliminary experiments designed to explore ribozyme-type cleavage of endogenous nucleic acids in normal or scrapie-infected brain were unsuccessful (not presented here).

In contrast, experiments in which total brain nucleic acid was incubated with dNTP precursors and reverse transcriptase (RT), in the absence of any primer, generated an alkali-sensitive band of around 300-350 nt (not shown); we infer that the labelled bands are likely to be partial extension products of 7S small cellular RNAs (sequences compiled in Shumyatsky & Reddy, 1993). Previous self-priming experiments (Suh et al., 1989) had centred on small RNAs 7SL, 7SK and 7-2/MRP; native 7SL failed to self-prime under their conditions but 7SK and 7-2 were reported to do so. 7SL (299 nt in this study) is an integral component of the signal recognition particle (Walter & Blobel, 1982) and may also play a role in ribosome biogenesis (Kiss et al., 1992). Neither 7SK (∼331 nt) nor 7-2 (∼275 nt) has been ascribed a definite function, though some work suggests that 7-2 RNA is involved in mitochrondrial RNA and/or tRNA processing (Chang & Clayton, 1989; Liu et al., 1994). Other candidate species selected include U3 and H1 (also known as 8-2). U3 is involved in early processing of ribosomal RNA transcripts (Hartshorne & Agabian, 1994) and may also be capable of self-priming reverse transcription (Bernstein et al.,1983), while H1/8-2 is a component of RNase P, an enzyme involved in the maturation of tRNA precursor molecules (Baer et al., 1990). We then set out to characterise the sequence and abundance of these small RNA species, in parallel, from scrapie-infected hamster brain and uninfected controls.

### Amplification of small RNA species from scrapie-infected and normal hamster brain

We pursued our studies using nucleic acid from hamster brain infected with 263K scrapie, a TSE strain that (on the basis of limiting dilution experiments) appears to contain one of the highest infectious titres (Kimberlin & Walker, 1977). For reverse trancription and PCR (RTPCR), oligonucleotide primers were designed corresponding to the candidate small RNA species and, in some cases, with 5’ extensions containing appropriate restriction sites to facilitate cloning. Primers were designed from known rat, mouse and/or human sequences (Methods). In general, the 3’ oligonucleotide was employed to prime reverse transcription (RT); subsequent PCR amplfication employed both oligonucleotides. In some cases random hexameric priming was also utilized (see below).

### RNAs 7-2, 7SL, U3, H1/8-2

For 7-2, using the 3’ primer for the RT step and both 5’ and 3’ primers for the PCR, a single product of approximately 270 nt was generated as expected and cloned. Four clones, two from normal hamster brain (61, 62) and two from scrapie-infected brain (32, 34) were selected and sequenced. Clones 61 and 32 (Figure 2) were highly homologous to mouse and rat 7-2 sequences. Clones 62 and 34 were novel sequences (discussed further, below). Direct sequencing of the 270 nt RTPCR product (uncloned) from normal and scrapie-infected brain was also performed; this failed to reveal any obvious differences between the nucleotide sequences (not shown).

**Figure 2.**
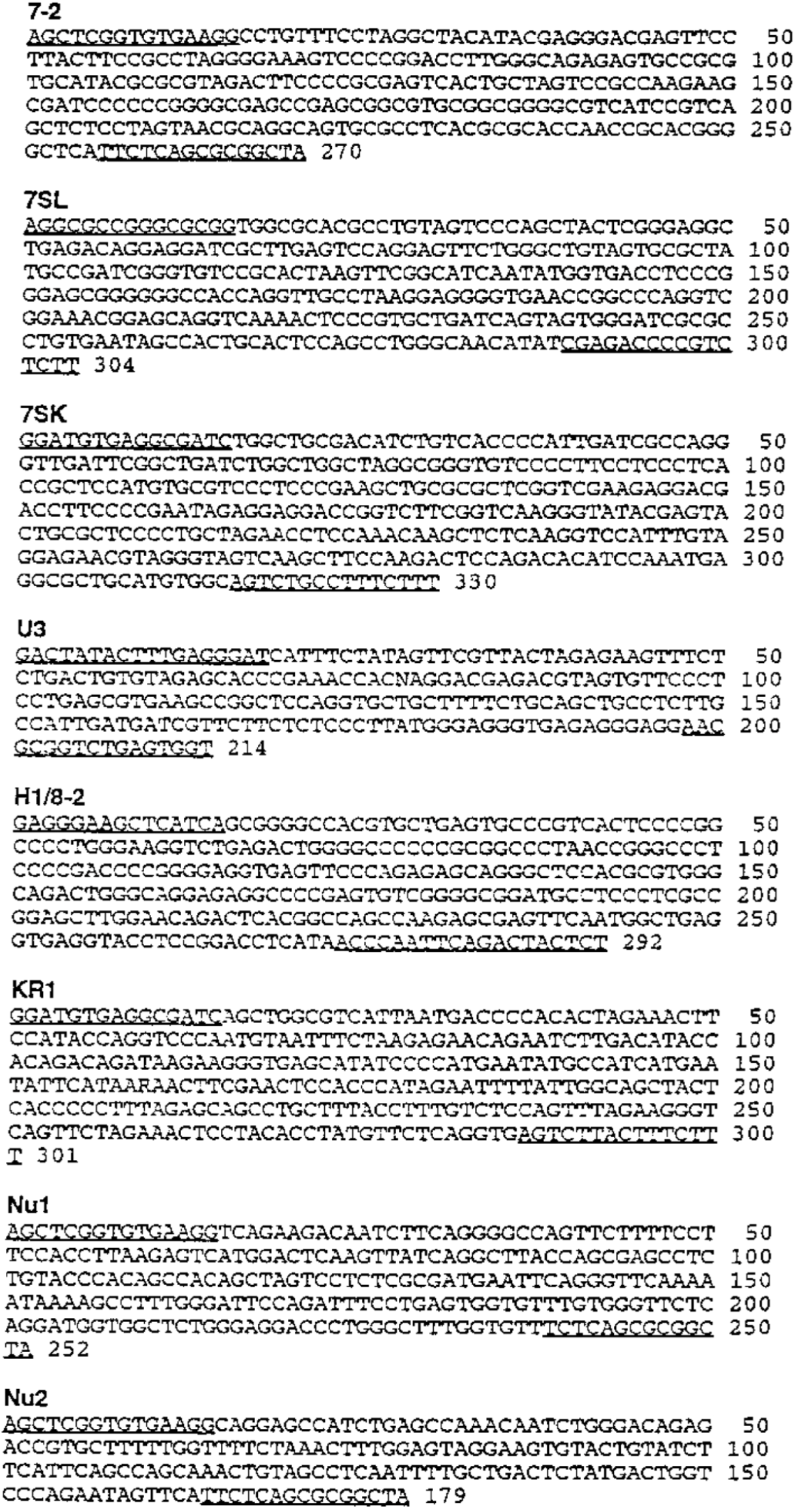
Sequences of small RNA species obtained from scrapie-infected hamster brain. Sequences corresponding to the PCR primers employed (lacking terminal additions of restriction sites) are underlined. These sequences have been submitted to the GenBank database under the accession numbers 00 (7-2); 00 (7SL), 00 (7SK), 00 (U3), 00 (KR1), 00 (Nu1), and 00 (Nu-2).

For RNA 7SL, random hexamers or the 3’ primer were included to facilitate RT copying of RNA into DNA; 7SL sequences were then PCR amplified using specific primers. A ∼300 nt product was generated from RNA from both normal and infected hamster brain. Other minor bands appeared to differ between normal and scrapie-infected brain but this was not reproducible. The major RTPCR product was cloned into plasmid vectors; independent clones from uninfected and infected brain were subjected to DNA sequence analysis (Figure 2). No differences were observed, arguing that the major species of 7SL in infected brain does not represent the genome of 263K scrapie.

For U3 and H1 the primary RTPCR products (∼210 and ∼290 nt) were subjected to direct sequencing without recourse to cloning. No differences in nucleotide sequence were noted between products from normal and scrapie-infected brain, arguing that these species are unlikely to be related to the disease agent. The sequences of hamster 7SL, 7-2, U3 and H1 are presented in Figure 2.

### Amplification with primers corresponding to 7-2 reveals new candidate species

Cloning of the 7-2 PCR product generated 2 plasmids containing smaller insert fragments (see above): clone 62 (from normal brain) and clone 34 (from infected brain) contained RTPCR products of 252 nt and 179 nt respectively. The two fragments were sequenced (Figure 2) and were found to be unrelated to each other, with little homology to 7-2, nor to any sequences in the database (as determined by standard searching; not presented), though it was not possible to address possible terminal homology to the 7-2 primers employed. Primers specific to these two novel sequences (provisionally termed RNAs nu1 and nu2) were used to confirm their presence in both normal and scrapie brain RNA samples. Direct sequencing of the nu2 RTPCR product was performed on material from normal and infected brain but failed to reveal differences in primary sequence. Technical problems precluded direct comparative sequencing of nu1 from normal and scrapie-infected brain due to the low abundance of the molecule and the presence of many non-specific products following RTPCR. However, inspection of several independent molecules cloned following RTPCR failed to reveal any obvious differences between normal and scrapie-infected brain (not shown). The nucleotide sequences of hamster nu1 and nu2 are presented in Figure 2.

### Northern analysis using probes specific for 7SL, 7-2, U3, H1, nu1 and nu2

To address the abundance of these molecules in scrapie-infected brain, total RNA preparations from representative infected and control brains were resolved by gel electrophoresis and, following transfer to membranes, hybridized to probes specific for these RNA species. No differences in the electrophoretic mobility or abundance were detected for RNAs 7SL, H1 or nu2 (not presented).

RNAs 7-2, U3 and nu1 were not readily detectable by this method, presumably due to their low abundance (not presented). Despite careful analysis, semi-quantitative PCR (performed by serial sampling at different times during the PCR reaction) of 7-2, U3 and nu1 failed to reveal any obvious differences in the abundance of these species (not shown).

### RNA 7SK

In initial experiments first-strand cDNA synthesis took advantage of the documented self-priming ability of 7SK (see earlier). PCR amplification using specific 5’ and 3’ primers then generated a band of ∼300 nt from uninfected hamster brain material. Surprisingly, in some experiments no reproducible PCR product was obtained from self-primed cDNA derived from scrapie-infected brain RNA, while the starting material appeared identical on gel electrophoresis and the same preparations efficiently generated 7SK product when cognate primers were used (Figure S1). We suspected the existence of a subtle structural alteration that could inhibit self-primed RTPCR amplification of 7SK from scrapie-infected hamster brain.

Failure to amplify 7SK from infected brain does not discriminate between a failure of primed RT or a failure to PCR amplify (due to sequence and/or structural differences). We therefore performed reactions in parallel using either self-priming or priming assisted with the specific 3’ primer for the RT step. A ∼300 nt PCR product was obtained from infected brain material when the 3’ primer was included at the RT step, while the self-priming reaction did not reproducibly generate a RTPCR product (Figure S1). This argued that a subtle alteration inhibiting RTPCR might affect self-priming of reverse transcription.

Northern analysis of 7SK revealed that this molecule was over-represented in scrapie-infected brain (Figure S2).

### Amplification using 7SK-primers also amplifies a novel K-related species, KR1

Following RTPCR, two putative 7SK clones obtained from normal brain material were sequenced: one species was highly homologous to published 7SK sequences and is presumed to be the hamster 7SK orthologue (Figure 2); the other is discussed below. Two clones obtained from infected brain material were also studied; no differences were observed between the nucleotide sequences of the major 7SK species in infected brain and that present in normal brain.

The other clone obtained from normal brain material was found to differ substantially from the 7SK species. The sequence of this molecule, referred to as K-related molecule 1 (KR1), is presented in Figure 2. Two KR1 clones then obtained from two samples of uninfected brain were compared with two independent KR1 clones obtained from scrapie-infected brain. This revealed a single nucleotide substitution present in each of two scrapie-infected brain samples that could potentially represent a disease-specific alteration.

### KR1 is present in brain but abundance does not correlate with disease

To determine whether KR1 is indeed present in hamster brain, and is not an RTPCR artefact, we designed new sets of internal RTPCR primer pairs on the basis of our KR1 sequence data. These routinely generated molecules of the expected size by RTPCR of normal hamster brain RNA (not shown), confirming that KR1 is an authentic hamster brain transcript.

Quantitative Northern analysis on brain nucleic acids from scrapie-infected and uninfected animals was then performed using a KR1-based probe. KR1-specific probes failed to give a signal with RNA from either normal and infected brain, suggesting that the level of expression of KR1 is below the limit of detection by this technique. We then performed quantitative RTPCR, using internal KR1-specific primers, to explore the distribution of KR1 in normal and infected brain. In 10 representative samples of infected and uninfected brain, the abundance of KR1 did not correlate with disease status (not presented).

### KR1 sequence alterations are unrelated to disease

To determine whether KR1 might harbour internal and disease-specific alterations, as suggested above, we sequentially cloned 1.3 kb *Pst*I and 1.5 kb *Pst*I-*EcoR*I fragments of hamster genomic DNA hybridizing to KR1 (not presented). Figure 3A presents the sequence of the hamster genomic locus encompassing the KR1 segment.

**Figure 3.**
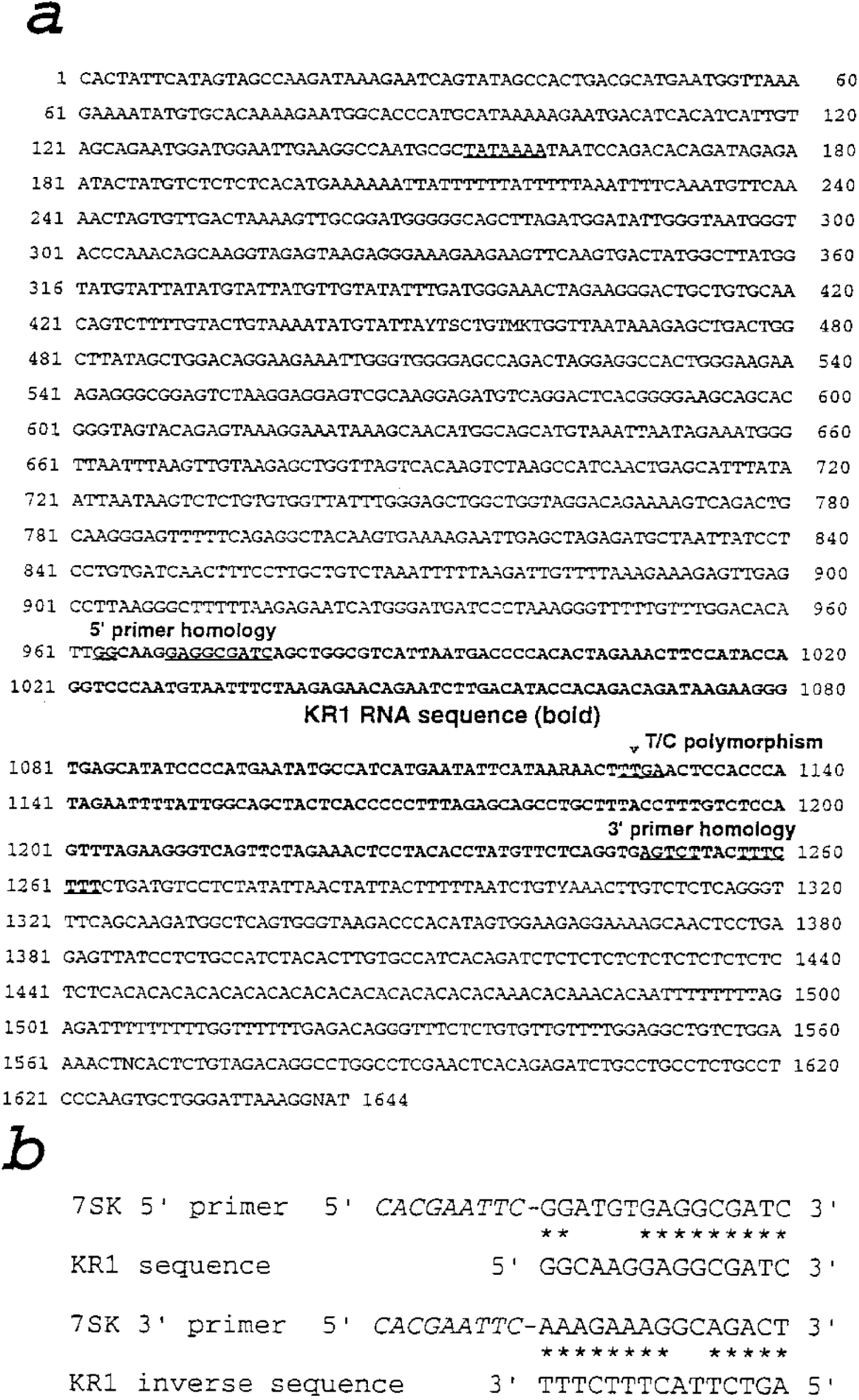
DNA sequence of the genomic locus encoding hamster KR1. (a): genomic sequence and regions corresponding to the 7SK primers employed; the T/C polymorphism (text for details) is indicated. (b): homologies between the 7SK primers and the KR1 sequence. The genomic sequence has been submitted to GenBank under the accession number 00.

Sequence analysis failed to reveal homologies in the GenBank/EMBL databases. Homologies between 7SK and KR1 were limited, except that the genomic KR1 sequence contains sequences similar to the 5’ and 3’ ends of the 7SK sequence against which the primers were designed (Figure 3B). To determine the size of the KR1 transcription product, total RNA was separated by gel electrophoresis, different size fractions were excised, and separately subjected to quantitative PCR. The majority of the KR1 primary transcript was found to be in a size fraction of 2000-4000 nt, substantially larger than 7SK (data not presented).

The genomic sequence (Figure 3A) was found to be identical to the RTPCR products, except that the TaqI site (TCGA) present in two independent RTPCR clones from scrapie-infected brain was replaced by TTGA in the genomic clone, corresponding to the sequence found in the RTPCR clone from uninfected brain. However, the TaqI sequence failed to correlate with disease, demonstrated by restriction analysis of RTPCR products obtained from a set of 10 infected or normal hamster brain samples (not shown); we surmise that the presence or absence of a TaqI site is probably a simple polymorphism within this hamster population.

## DISCUSSION

We have begun to explore the possible involvement of small nucleic acids in the etiology of the transmissible encephalopathies. These studies were encouraged by the diversity of different scrapie strains, incompatible with a protein-only hypothesis (introduction). We contend that mutation and/or recombination of endogenous RNA molecule(s) may give rise to a species capable of replication (introduction), exploiting *in vivo* processes such as RNA-dependent RNA polymerization and ribozyme-type cleavage. The hypothesis could potentially explain the enigmatic nature of the TSEs, that they may arise spontaneously (sporadic disease) but are subsequently transmissible by inoculation. While the incidence/onset of disease is clearly modulated by mutations or polymorphisms in a cellular gene (*PRN-P* in human) encoding the cell surface protein PrP, these observations would also be consistent with a role for PrP as a cellular receptor mediating internalization of the agent. Further, the finding that mice lacking a functional *Prn-p* gene fail to develop disease following inoculation of agent (Büeler et al., 1993) is consistent with this interpretation.

Many small RNA species present in rodent brain. Some of these, particularly those in the 7S group, have the capacity to self-prime reverse transcription. Accordingly we studied a selection of the most promising candidate RNAs. This was done by PCR amplifying and sequencing cDNAs from scrapie-infected (263K) hamster brains and from uninfected controls. Intriguing variations in our ability to PCR amplify the 7SK molecule from scrapie-infected brain, and its overabundance therein, led us to focus on this molecule, and a molecule termed K-related molecule 1 (KR1). Through intensive analysis, including cloning and sequencing of the genomic locus encoding KR1, we now argue that the major species of 7SK and KR1 are probably unrelated to disease status. To date our analysis has revealed no systematic differences between the abundance and/or sequence of any of the 8 candidate molecules investigated in normal and scrapie-infected hamster brain (Table 1).

**Table 1.** Candidate molecular species examined.

| RNA species<br>(nt)* | Length | Role, homologies | Reference |
| --- | --- | --- | --- |
| 7-2 (MRP) | 270 | mitochondrial RNA and/or tRNA processing | (Chang & Clayton, 1989) |
| 7SL | 304 | signal recognition particle; homology to Alu repeat elements | (Walter & Blobel, 1982) |
| 7SK | 330 | unknown | (Reddy et al., 1984) |
| U3 | 214 | ribosomal RNA processing | (Hartshorne & Agabian, 1994) |
| H1/8-2 | 292 | component of RNase P; involved in tRNA precursor maturation | (Baer et al., 1990) |
| KR1 | 301 | unknown, novel | this work |
| nu1 | 252 | unknown, novel | this work |
| nu2 | 179 | unknown, novel | this work |

\*Sizes of PCR products (nt, nucleotides); the true extents of the primary transcription product(s) in the hamster are unknown.

This work, however, has many limitations. First, mutations that lie under the sites complementary to the primer pairs or outwith the segment amplified will not be detected. Second, this analysis would probably not detect a modified RNA species which, in terminal brain, is substantially less abundant than the parent species. This is a major concern, because replication of the presumed agent in only a small subset of neurones could be sufficient to cause death of the host.

Third, putative modified and replication-competent species, particularly those propagated through many serial passages in a different host (such as 263K), could be so substantially modified that binding sites for the PCR primers may have been lost, in which case PCR amplification would fail to identify such species whatever their abundance. While not without biological risk, studies on sporadic cases of CJD by the approach described here could prove more fruitful.

Fourth, if the infectious agent is generated by rare recombination between two endogenous nucleic acids, primers designed from any one species would fail to PCR amplify the hybrid molecule. Fifth, molecular species which are refractory to RTPCR, for instance by covalent linkage of protein or other modifications that block copying by replicative enzymes *in vitro* (but not *in vivo*) would not be detected by this analysis. Sixth, species that cannot be propagated in *E. coli* (for instance because they cause lethality or are incompatible with plasmid replication) would not be detected in some of our analyses.

Although the present work fails to provide positive substantiation for the hypothesis, we believe that the contention that the etiologic agent arises from a modified endogenous nucleic acid has not yet been disproved. Future studies towards the identification of disease-specific nucleic acids will involve the use of more sophisticated technologies.

## ACKNOWLEDGEMENTS

This work was supported by grant funding from the Ministry of Agriculture, Fisheries and Foods (MAFF), by BBSRC support to the Centre for Genome Research, and by BBSRC and MRC support to the Neuropathogenesis Unit, Institute of Animal Health.

## NOTE

This MS was originally prepared in 1998 but was not published.

## Supplementary Data

**Figure S1.**
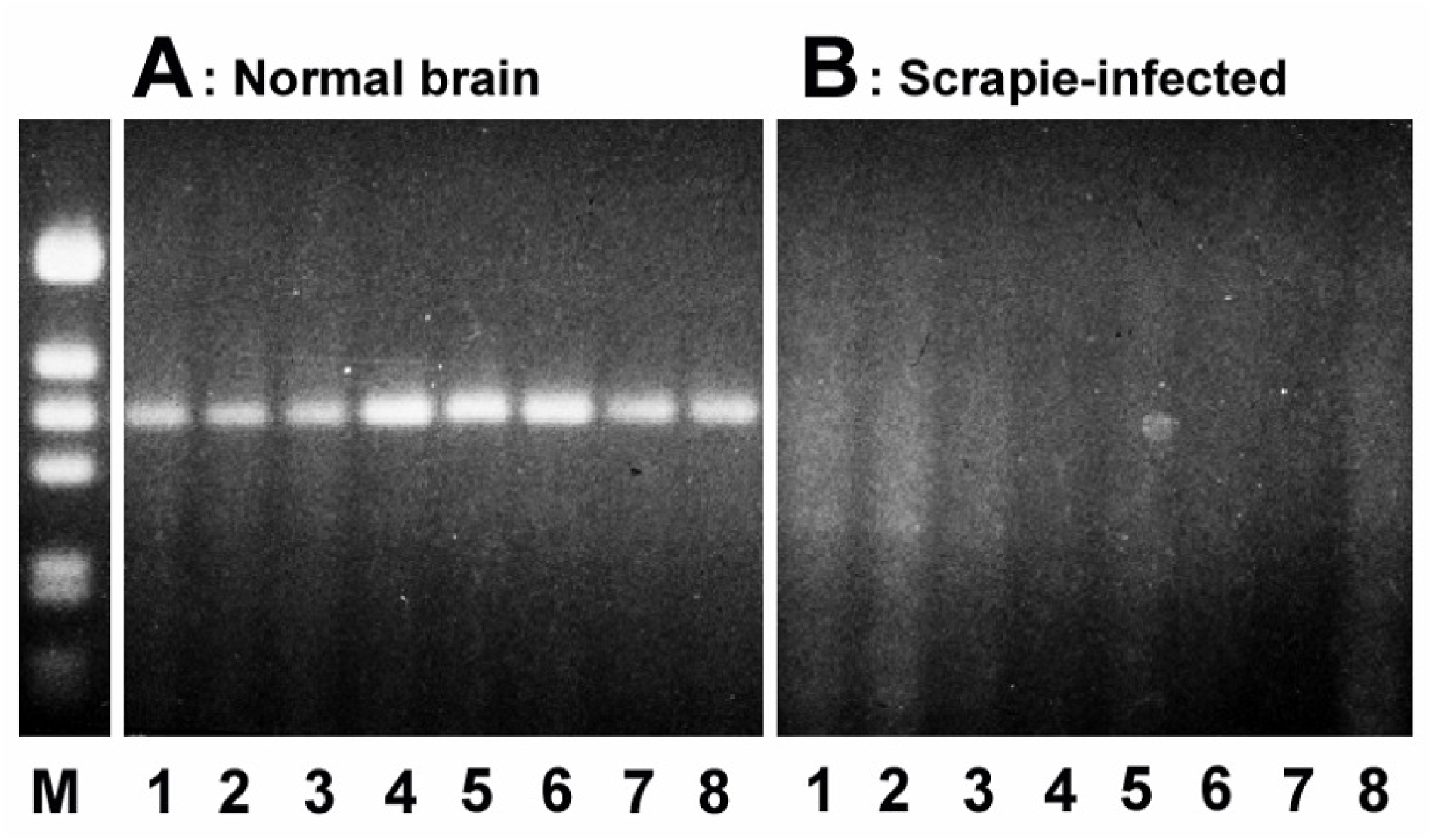
Self-priming of reverse transcription (RT). RNA 7SK is known to self-prime reverse transcription (RT). Extracts of control brain (A) and strain 263K brain (B) were incubated with RT and dNTPs, and resolved by agarose gel electrophoresis. No product was observed in scrapie-infected brain.

**Figure S2.**
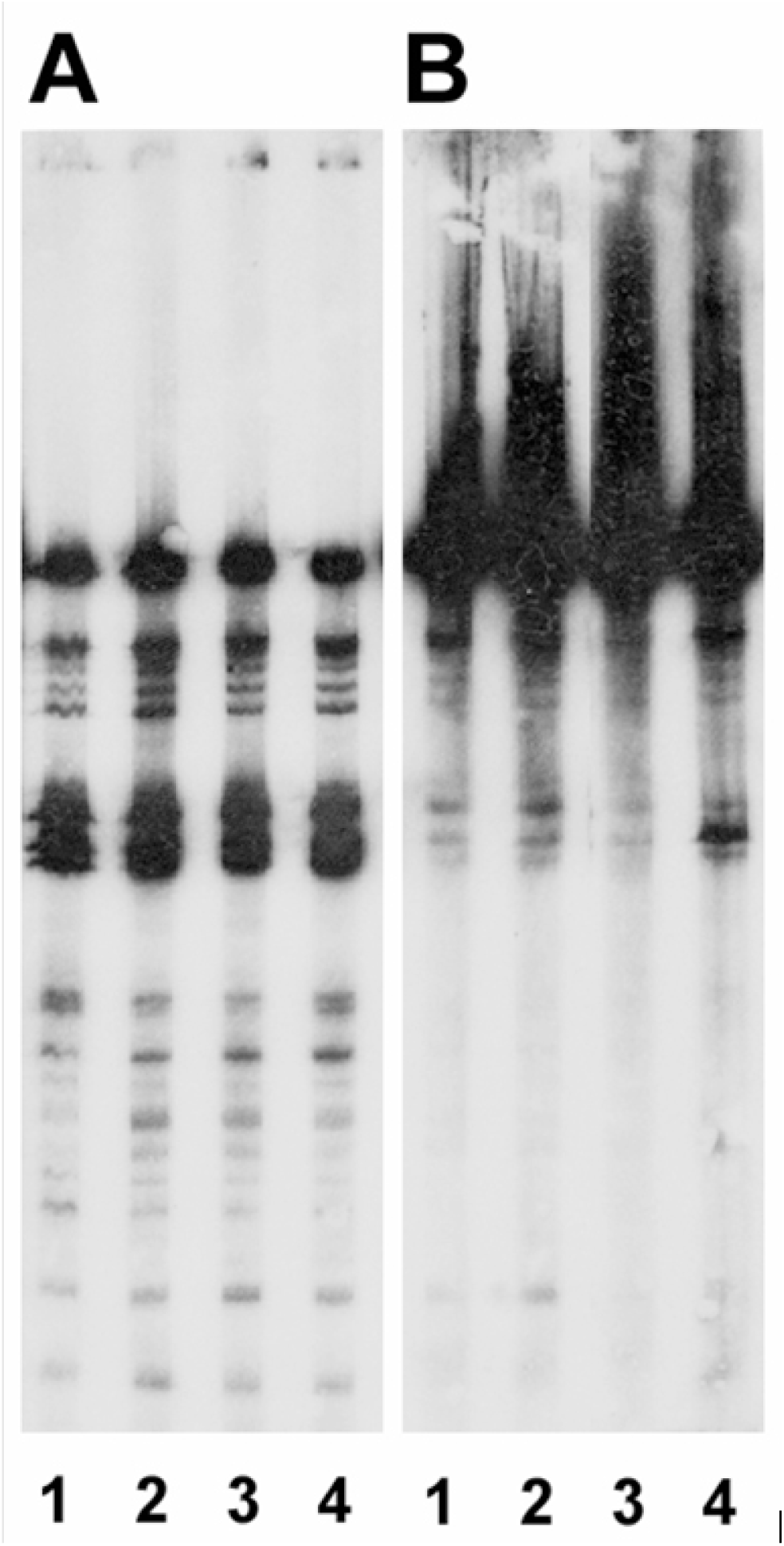
Abundance of 7SK in control (A) and scrapie-infected (B) brain assessed by northern blotting. Image J scanning indicates a 20–50 fold increase in abundance of 7SK in scrapie infection.

